# BioClaw: Human-Bot Research Collaboration Ecosystems in Group Chats

**DOI:** 10.64898/2026.04.11.716807

**Authors:** Mingyang Xu, Jiaxian Yan, Runchuan Feng, Qingran Cai, Peng Zhang, Chuang Zhao, Cao He, Zongting Wei, Jianping Li, Shiyi Lin, Hongyu Dong, Ruofan Jin, Tingjun Hou, Qi Liu, Zaixi Zhang

## Abstract

Day-to-day research discussions in group chats often generate hypotheses, analysis requests, and interpretation decisions, yet executing those analyses still requires researchers to leave the conversation and rely on fragmented local tools, databases, visualization software, and literature search engines. In this work, we present **BioClaw**, a human-bot research collaboration ecosystem that converts natural-language requests in group conversations into tool-grounded analyses executed within isolated Docker containers. Deployed across 8 messaging platforms, BioClaw turns each group chat into a persistent execution workspace. Its design combines multi-channel orchestration, per-group state and workspace management, and isolated containerized execution for reliable shared use over long-lived conversations. To support practical research workflows, BioClaw combines containerized execution with preinstalled 31 biomedical tools and 95+ skills. The application of BioClaw spans various biomedical domains (e.g., genomics, clinics, structural biology) and data modalities (e.g., sequencing data, EHR data, protein structure data). These results establish the viability of embedding executable, tool-rich agent workflows within shared digital workspaces, positioning group chats as a transformative paradigm for collaborative scientific discovery and innovation.

## 1. Introduction

Daily discussions on messaging platforms such as WhatsApp, Slack, and WeChat have become part of how research and clinical teams communicate, coordinate, and exchange questions (Carmona et al., 2018; Gofine and Clark, 2017; Nature Methods, 2020). Yet turning those discussions into executable bioinformatics analyses and bringing the resulting outputs back into the same conversational setting remains difficult in practice (Suetake et al., 2024; Zhang and Cranshaw, 2018). The challenge arises not only from the intrinsic complexity of biological data analysis, which often requires multiple computational tools, but also from the fragmented and evolving software ecosystem on which such workflows depend (Suetake et al., 2024). A central challenge, therefore, is how to connect conversational requests with reliable analytical execution and return the resulting outputs to the same collaborative context, thereby making chat-based scientific collaboration more actionable in practice (Galaxy Community, 2024; Zhang and Cranshaw, 2018).

Relevant prior work has made progress toward this goal. LLM-based agent frameworks (Anthropic, 2026; Jin et al., 2025; Schick et al., 2023; Yao et al., 2023) have shown that complex, multi-step tool use can be orchestrated directly from natural-language instructions. Workflow platforms such as Galaxy (Galaxy Community, 2024) reduce integration friction by providing a structured interface for composing and executing bioinformatics pipelines. Despite these advances, an important gap remains at the intersection of conversational agents and bioinformatics workflows. Existing approaches have not fully addressed how executable bioinformatics capabilities can be embedded into long-lived group chats, where ideas are generated and results are discussed over time.

To bridge this gap, we present **BioClaw**, a human-bot research collaboration ecosystem in group chats. Deployed across eight major messaging platforms, including WhatsApp, Feishu, WeCom, Discord, Slack, WeChat, QQ, and a local web UI, BioClaw converts natural-language requests from long-lived group conversations into executable, tool-grounded workflows. These workflows run inside isolated containers and return results directly to the same shared thread. Rather than treating chat as a thin interface for one-shot question answering, BioClaw treats each group as a persistent computational workspace in which conversational context, uploaded files, intermediate artifacts, and analysis results can accumulate and be reused across turns. This design allows scientific discussion, analytical execution, and result interpretation to remain co-located within the team’s existing communication channel.

At the system level, BioClaw combines a multi-channel orchestration layer with a prebuilt bioinformatics runtime. The orchestration layer normalizes requests from heterogeneous chat interfaces while maintaining per-group state, workspace isolation, and execution queues, enabling concurrent use across multiple groups and platforms without losing contextual continuity. The runtime layer packages bioinformatics command-line tools, Python libraries, and reusable skills, enabling practical bioinformatics analyses spans various domainssuch as sequence similarity search, sequencing quality control, differential-expression visualization, protein structure rendering, and literature retrieval.

To validate the practical utility of BioClaw, we deployed it across eight messaging platforms and conducted a series of representative application case studies. These studies examine whether a chat-native agent can accept diverse scientific inputs, invoke appropriate tools or skills, and return useful artifacts within the same shared conversational context (Sections 3 and 4). Our results indicate that BioClaw can serve as a practical foundation for integrating bioinformatics analysis into long-lived group chats, providing a shared cross-platform execution substrate for collaborative scientific discovery and innovation.

In summary, this report makes four main contributions:

- We present **BioClaw**, a human–bot research collaboration ecosystem in group chats that embeds executable bioinformatics workflows directly into long-lived group conversations, turning the chat thread itself into a persistent computational workspace rather than a front end to isolated analyses.
- We design and implement a practical system architecture for reliable multi-group operation, combining multi-channel message orchestration, per-group state and workspace management, isolated containerized execution, dual backend support, and secure host–container communication mechanisms.
- We build a reproducible bioinformatics runtime that integrates a broad collection of preinstalled command-line tools, Python libraries, and reusable skills, enabling diverse analysis tasks to be invoked from natural-language requests without requiring users to leave the conversation environment.
- We validate BioClaw through real-world deployment across eight messaging platforms and a series of representative application case studies, showing that the system is both practically deployable in heterogeneous chat environments and effective in supporting diverse bioinformatics workflows **within** the same collaborative thread.

## 2. The BioClaw System

### 2.1. Design Objective

BioClaw is designed around a single objective: to make the group chat thread itself the execution locus of a bioinformatics research workflow, rather than a front end to one-shot question answering. In laboratory settings, requests rarely end at a single textual reply; they involve uploaded files, parameter refinement, multi-step tool execution, and the return of interpretable artifacts such as tables, plots, rendered structures, and derived files.

Formally, for each group *g*, the system maintains a persistent context

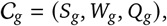

where *S*_*g*_ denotes conversational state, *W*_*g*_ the group workspace, and *Q*_*g*_ the execution queue. Given a new message m, in group *g*, BioClaw maps the pair (*m*_*t*_, *C*_*g*_) to an executable action sequence *a*_*t*_, returns artifacts *o*_*t*_, and updates the context:

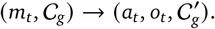

This formulation makes two system requirements explicit. First, interaction must be *persistent and group-centric:* bioinformatics work is negotiated across participants and across turns, so each group receives its own session context, workspace, and execution queue. Second, capability must be *too/-grounded:* the model plans and orchestrates, but substantive analysis is carried out through external software, Python libraries, and domain-specific skills (Jin et al., 2025).

### 2.2. System Overview

At a high level, BioClaw implements a group-scoped execution loop (Figure 2). A request arrives through one of the supported chat interfaces and is normalized by the orchestrator, which resolves the corresponding group state and workspace. The request is then dispatched to a containerized runtime where the agent invokes bioinformatics tools or reusable skills to perform the analysis. The resulting text, files, or images are returned to the same conversation thread and become part of the context for subsequent turns. Details about a request lifecycle can be found in Figure. S1.

**Figure 1.**
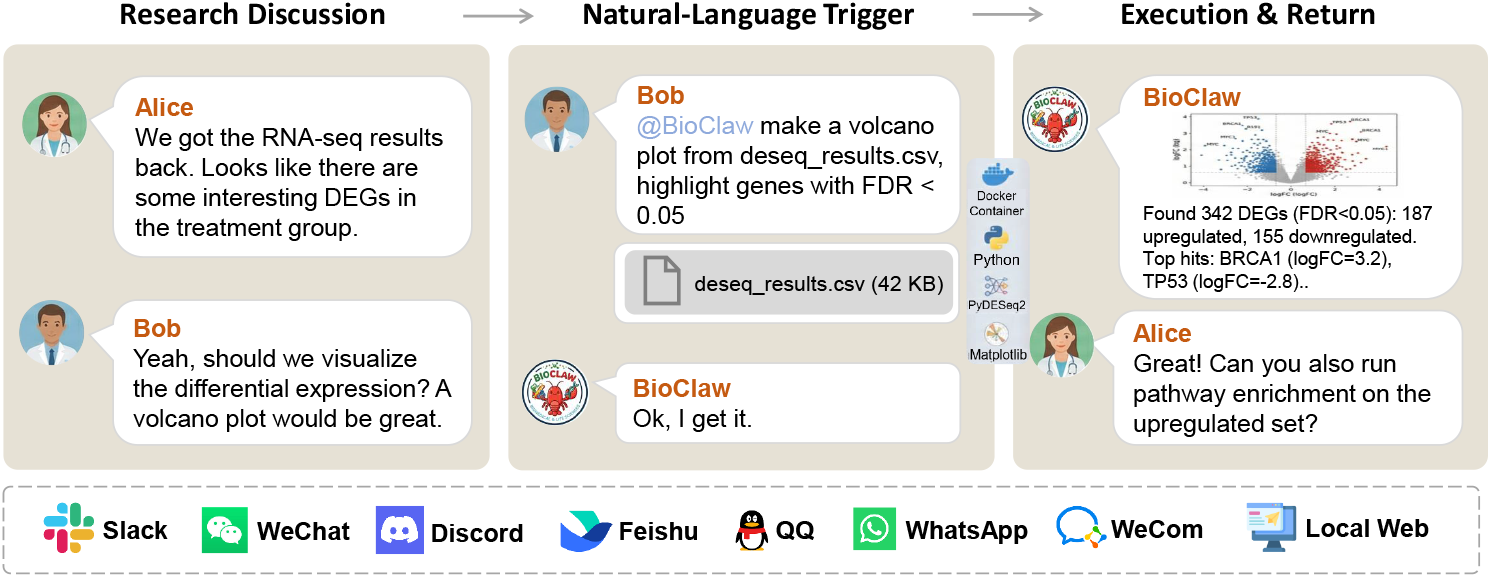
Human–bot research collaboration in BioClaw. Researchers discuss experimental results in a group chat *(left)*, issue natural-language requests to trigger analysis *(center)*, and receive executable, tool-grounded outputs such as figures, structured summaries, and follow-up suggestions within the same conversation thread *(right)*. This workflow illustrates how conversational interactions are transformed into persistent, executable scientific processes across eight messaging platforms (bottom).

**Figure 2.**
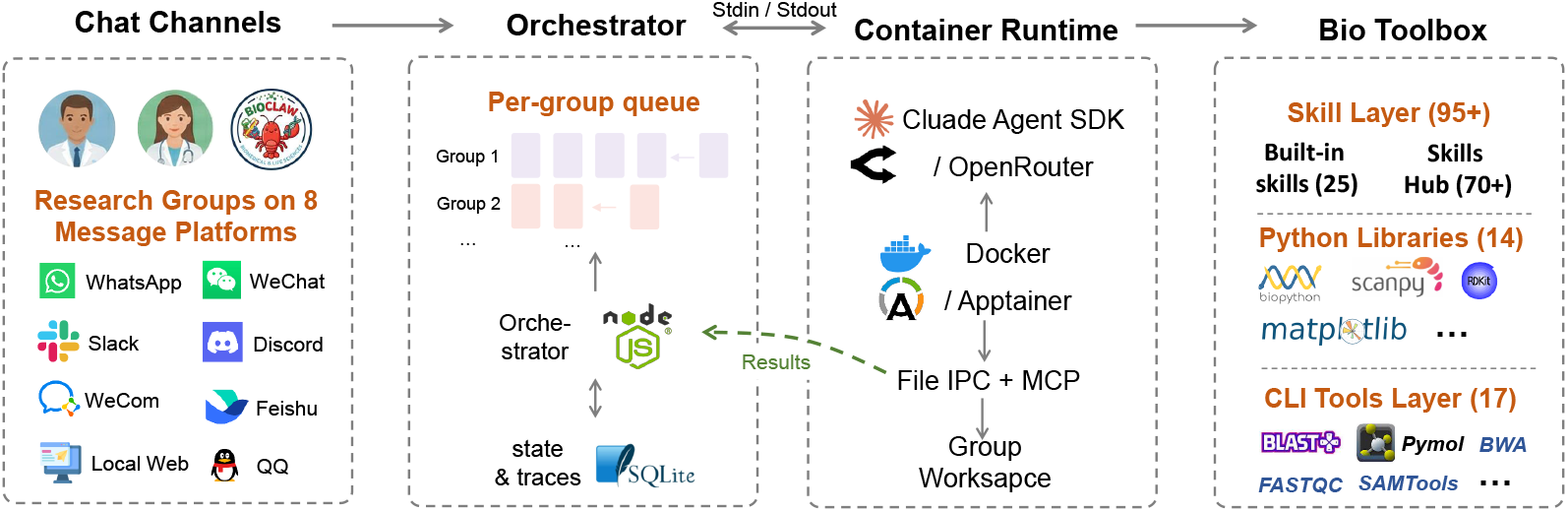
User requests from eight chat channels are routed through a Node.js orchestrator with per-group queueing and SQLite-based state management, then executed in isolated containers. Each container supports dual model backends, workspace access through IPC/MCP, and a bioinformatics runtime composed of reusable skills, Python libraries, and command-line tools, enabling natural-language group-chat requests to be turned into executable analyses.

The remainder of this section describes the two critical modules that support this loop: the orchestration infrastructure inherited from NanoClaw (QwibitAI, 2026) (Section 2.3), and the bioinformatics runtime—comprising preinstalled tools, libraries, and domain—specific skills—that constitutes BioClaw’s bioinformatics capabilities (Section 2.5).

### 2.3. Orchestration Infrastructure

BioClaw inherits its orchestration layer from NanoClaw (QwibitAI, 2026), where each chat platform is wrapped by a *channel adapter* that normalizes platform-specific events into a unified message representation and persists them in SQLite. Any subset of the eight supported interfaces—WhatsApp, Feishu, WeCom, Discord, Slack, WeChat, QQ, and a local web UI—can be enabled independently while the orchestration logic and execution runtime remain shared. The *group* is the primary computational unit: each group maintains a dedicated workspace, session identifier, message cursor, and execution queue, with cursor advance-then-rollback semantics ensuring transparent failure recovery under concurrent multi-group operation.

The execution layer is designed to remain agnostic to both model provider and container run-time. Within the same containerized environment, BioClaw supports two model backend paths. The Anthropic path uses the Claude Agent SDK directly, exposing its native tool suite and MCP integration; the OpenAI-compatible path targets providers exposing a `/chat/completions` interface and implements an explicit tool-calling loop with a compact tool set, including shell execution, file I/O, and artifact return primitives. To close the capability gap between the two paths, `read_file, write_file`, and `list_files` are provided for OpenAI-compatible models, allowing them to **in**spect workspace contents and load full skill definitions rather than relying only on skill summaries. Backend selection is resolved at container startup from environment variables and remains transparent to the orchestrator, as both paths emit results through the same stdout-based IPC protocol.

To broaden deployment beyond standard servers, BioClaw further abstracts over the container runtime itself (as shown in Figure. S2). While Docker is the default backend, the system also supports Apptainer for HPC environments where Docker is often unavailable. The orchestrator delegates runtime-specific lifecycle operations—including launch, shutdown, and orphan cleanup—to a backend module selected through the `CONTAINER_RUNTIME` setting. This abstraction leaves the agent loop, skill system, and IPC design unchanged: the same runtime environment is reused across backends, differing only in packaging format and invocation mechanics.

Every analysis runs inside a Docker container that provides filesystem isolation, a reproducible environment, and a controlled security boundary. Host–container communication follows a dual-channel IPC design: stdin/stdout for prompts and streaming output, and shared filesystem directories for follow-up messages, artifacts, and scheduling. The agent accesses structured tools, including `send_message, send_image`, and scheduling primitives, through an MCP (Anthropic, 2024) server.

### 2.4. Reproducible notebook generation

To bridge the gap between conversational interaction and reproducible analysis, BioClaw automatically generates Jupyter notebooks from each successful agent run. During container execution, both model backends emit structured trace events, including agent reasoning steps and tool invocations, through the IPC layer. Upon run completion, the orchestrator reconstructs a chronologically ordered notebook from these events: Python code executed via shell heredocs is extracted into executable code cells, file-write operations are rendered with their content, and agent reasoning is preserved as interleaved Markdown cells. The resulting `.ipynb` file is saved to the group workspace and conforms to the standard nbformat `v4` specification, making it immediately executable in JupyterLab or VS Code. This design ensures that every chat-initiated analysis produces a self-contained, re-runnable artifact without requiring users to manually transcribe or export their work.

### 2.5. Bioinforrnatics Runtime

While the orchestration infrastructure is domain-agnostic, BioClaw’s domain-specific capabilities lies in the bioinformatics runtime bundled inside each container. Instead of installing software on demand, BioClaw packages a curated and reproducible environment directly into the image, so that every execution starts from the same analysis-ready state.

As shown in Figure. 3, the runtime includes 17 command-line tools for sequence search, read alignment, BAM/VCF processing, quality control, transcript quantification, and molecular visualization, together with 14 Python libraries for sequence I/O, numerical computing, visualization, single-cell analysis, differential expression, and cheminformatics. Consistent with the isolation model in Section 2.3, all processes run as a non-root user (`node`), with /`workspace/group, /workspace/global`, and /`workspace/ipc` serving persistent storage, shared read-only resources, and host–container communication, respectively. The image also includes Chromium and CJK fonts for browser automation and multilingual figure rendering, and initializes in under ten seconds on typical hardware.

**Figure 3.**
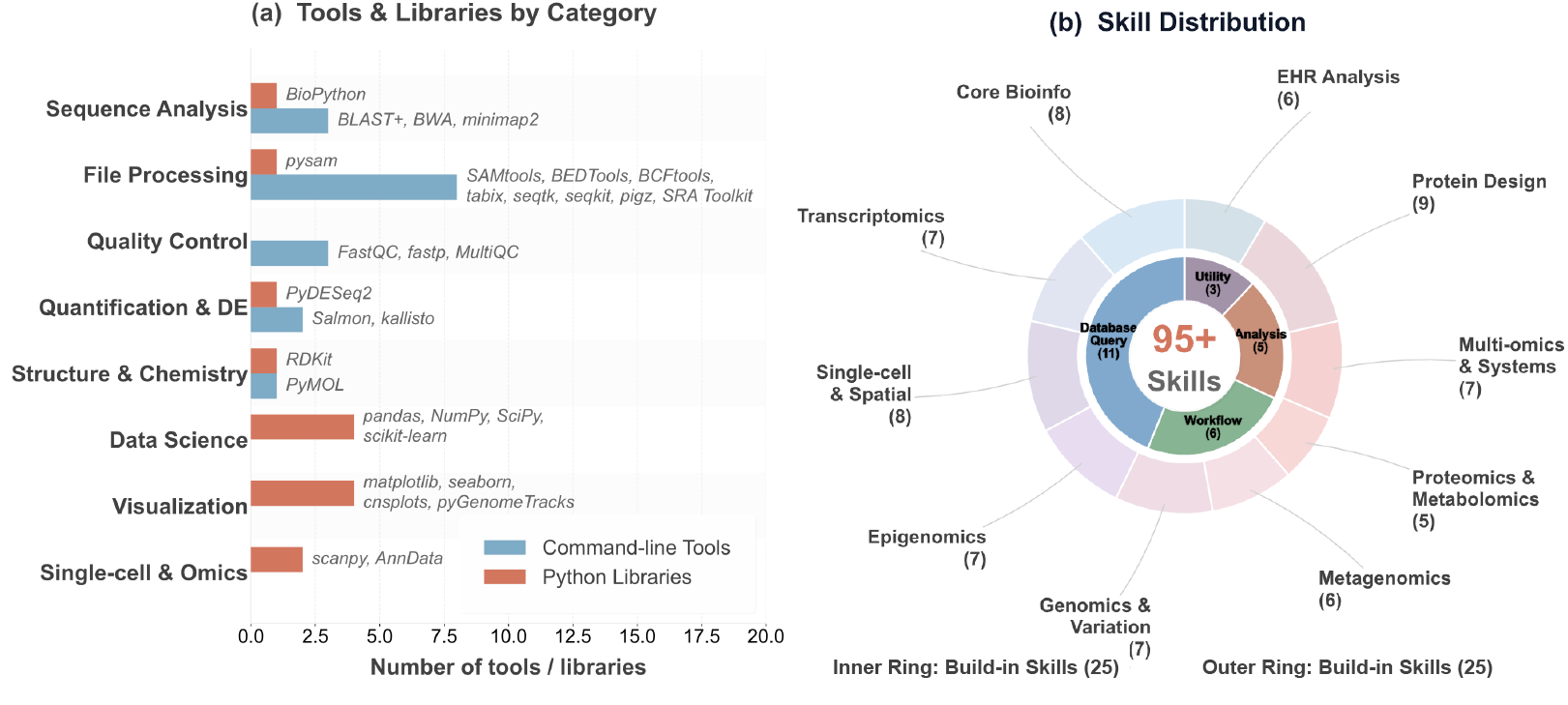
Bioinformatics runtime composition. **(a)** Distribution of 17 command-line tools and 14 Python libraries across eight functional categories. **(b)** Nested distribution of 95+ skills: inner ring shows 25 built-in skills in four categories; outer ring shows 70+ community-contributed Skills Hub entries across ten domains.

Beyond preinstalled software, BioClaw provides a two-tier skill architecture for agents (Figure. 3 and S2). As shown in the figures, there are 25 built-in skills packaged in the image and loaded at startup, covering common workflows such as BLAST search, differential expression analysis, single-cell RNA-seq preprocessing, database access, literature retrieval, and structure rendering. Besides, to extend coverage without enlarging the base image, BioClaw also Skills Hub skills (70+ across 10 domains) from a community-maintained repository^1^, letting the agent discover relevant entries and execute them on demand.

### 2.6. Skill Design Philosophy

In BioClaw, Skills are designed not merely as thin wrappers around individual tools, but as workflow-level abstractions for recurrent research tasks. This distinction is particularly important in long-lived group chats, where users may interrupt an analysis, inspect intermediate results, refine requirements, and resume work across multiple conversational turns. Under such conditions, a useful Skill must support not only execution, but also inspection, recovery, and collaboration.

In practice, we design Skills around three properties (as shown in Figure. S4). First, they are *modular and composable:* each Skill encapsulates a semantically coherent subtask and exposes outputs **in** forms that can be consumed by later steps or by other Skills. This allows a complex research workflow to be decomposed into checkpointed stages rather than executed as an opaque monolithic procedure. Second, they are *artifact-persistent:* intermediate outputs are written to the group workspace instead of being retained only in transient model state. This makes intermediate products inspectable by users, supports partial reruns, and provides an explicit chain of provenance across stages. Third, they support *human-in-the-loop control:* users can review intermediate results, provide corrective feedback, and redirect the workflow at appropriate checkpoints without discarding all prior progress. These properties extend the role of Skills beyond tool access alone.

## 3. Representative Applications

To illustrate the practical utility of BioClaw, we present a set of representative application cases that span core bioinformatics workflows and input modalities. Rather than evaluating the system as a conventional question-answering agent, we focus on whether it can accept heterogeneous scientific requests issued in natural language, invoke the appropriate tools or skills, and return interpretable results to the same shared conversation context. Figure 4 summarizes eight such cases, covering sequencing quality control, sequence similarity search, differential-expression visualization, structural biology, literature retrieval, protein structure prediction, binding-site inspection, and wet-lab image review.

**Figure 4.**
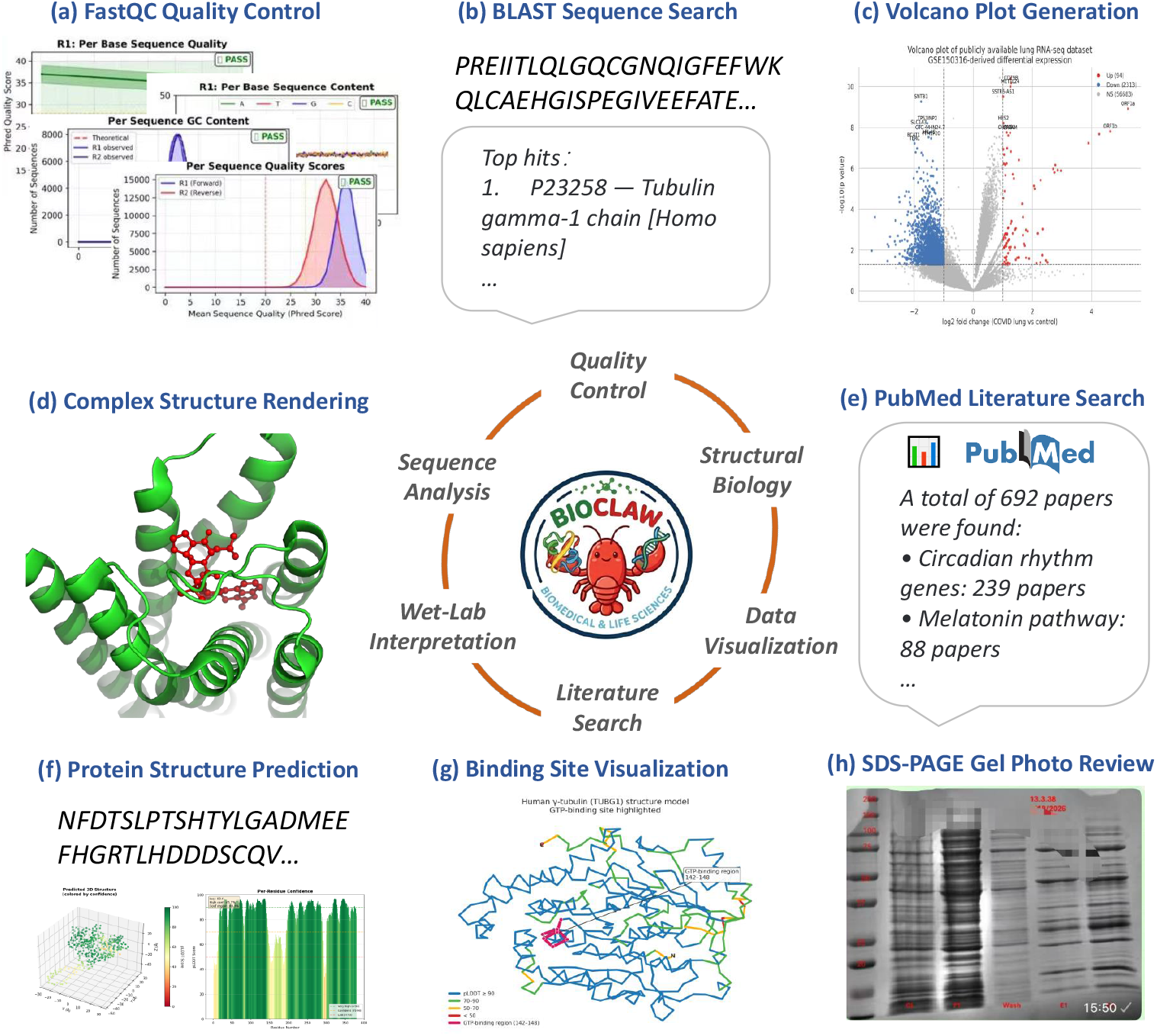
Representative BioClaw demos. The 8 example panels show (a) FastQC quality control for paired-end reads, (b) BLAST sequence search with structured top-hit summaries, (c) volcano plot generation from differential-expression data, (d) PyMOL-based protein structure rendering, (e) PubMed literature search and structured summarization, (f) Protein structure prediction with ESMFold, (g) binding-site visualization around ligand AQ4, and (h) SDS-PAGE gel photo review from a chat-uploaded image.

### 3.1. Representative Case Studies

The examples in Figure 4 are intended to illustrate the breadth of workflows that can be handled within the same chat-native execution framework. These cases involve markedly different input types, including uploaded FASTQ and CSV files, pasted nucleotide or protein sequences, PDB structures, and experimental images. Despite this heterogeneity, the user interaction pattern remains uniform: a natural-language request is issued in the group conversation, executed through the containerized runtime, and the resulting artifacts are returned to the same thread for inspection and follow-up discussion.

Panels (a)–(c) demonstrate file-centric and sequence-centric analytical workflows. The FastQC example shows that BioClaw can process raw sequencing reads and return quality-control summaries directly in chat. The BLAST case illustrates database-backed sequence similarity search with structured reporting of top hits. The volcano-plot example further shows that tabular differential-expression results can be converted into publication-style visual summaries without requiring users to switch to external analysis software.

Panels (d), (f), and (g) highlight structure-oriented molecular analysis. Together, these examples span complementary stages of structure-aware bioinformatics workflows, including global protein rendering, de novo protein structure prediction with ESMFold, and local binding-site visualization around ligand AQ4. They are important because they show that the same chat-native framework can support both the generation of structural hypotheses from sequence and the downstream inspection of three-dimensional molecular context. This reflects a common collaborative pattern in bioinformatics practice: a conversation may begin with a sequence or structural artifact and then iteratively evolve toward increasingly specific structural questions.

Panels (e) and (h) extend the system beyond standard command-line bioinformatics. In panel (e), the agent performs literature retrieval from PubMed and returns structured summaries grounded in retrieved papers. In panel (h), the workflow operates on an experimental gel image, showing that the same execution framework can support image-based wet-lab interpretation in addition to dry-lab computational analysis.

Taken together, these cases show that BioClaw supports a broad range of bioinformatics workflows under a consistent interaction model. Across all examples, users issue requests in natural language, the system invokes the required tools, and the resulting artifacts are returned directly into the same conversation thread. These artifacts include quality-control reports, ranked search results, statistical plots, molecular renderings, predicted structures, and image-based assessments. More importantly, these results demonstrate that BioClaw bridges the gap between conversational intent and executable scientific workflows, enabling heterogeneous data modalities and analysis tasks to be handled within a unified, chat-native computational environment.

### 3.2. Long-horizon Manuscript-planning

Beyond the task-specific examples in Figure 4, BioClaw can also support longer-horizon research workflows through workflow-oriented Skills. A representative example is a manuscript-planning Skill that organizes an early-stage research idea into a structured package for downstream discussion, implementation, and validation within the same group-chat workspace (Figure. S7).

In a typical use scenario, a user provides a research topic together with partial context such as related papers, baseline methods, intended innovations, candidate tasks, datasets, and evaluation criteria. The Skill then decomposes the request into multiple stages, including related-work retrieval, novelty assessment, task and dataset specification, metric design, analysis planning, figure planning, and draft generation. Rather than returning only a final text response, the workflow materializes intermediate artifacts at each stage as files in the group workspace. These artifacts can be inspected directly by users and reused as inputs to downstream stages, allowing the workflow to remain transparent and resumable across turns.

A key feature of this process is that user feedback can be incorporated at the level of workflow checkpoints rather than only at the final output. For example, if a user judges that a proposed figure or task design is inadequate, the workflow can return to the corresponding earlier stage, revise the affected design, and continue from there without rerunning unrelated stages. In this way, the Skill preserves efficiency while keeping scientific judgment with the user. The resulting workspace may include structured outputs such as novelty assessments, task definitions, dataset catalogs, metric systems, figure designs, manuscript drafts, refinement logs, and implementation plans, all of which remain available for later review and continuation.

### 3.3. Human-in-the-loop Research Assistant

Beyond directly completing diverse bioinformatics tasks such as those illustrated above, BioClaw is also useful in how it supports the user throughout the analytical process. Rather than serving only as a conversational front end, it can function as a practical human-in-the-loop research assistant within the same shared chat context.

Without such a system, many routine tasks would require users to identify appropriate resources, switch across multiple websites or software tools, prepare and upload data, write commands or analysis scripts, run external programs, and manually organize intermediate outputs for interpretation. With BioClaw, substantial parts of this process can often be coordinated within one or a few prompts, reducing setup overhead, tool switching, and manual orchestration.

Importantly, this does not remove the user from the workflow. Instead, the system assists with execution, tool use, and artifact generation, while the user remains responsible for defining the task, inspecting the returned results, and making scientific judgments. In this sense, BioClaw helps researchers focus more on interpretation and iteration, while still remaining useful at the conversational level even when a request falls outside the scope of the currently available bioinformatics tools.

## 4. Practical Deployment and Accessibility

Beyond demonstrating representative bioinformatics workflows, an important contribution of BioClaw lies in its practical deployment and accessibility. Rather than limiting the system to a single prototype interface, we implemented and deployed BioClaw across eight messaging environments, including WhatsApp, Feishu (Lark), WeCom, Discord, Slack, WeChat, QQ, and a local web UI, as shown in Figure 5. This broad platform coverage shows that the system is not merely a lab-side proof of concept, but a deployable chat-native bioinformatics assistant that can be integrated into heterogeneous real-world communication settings.

**Figure 5.**
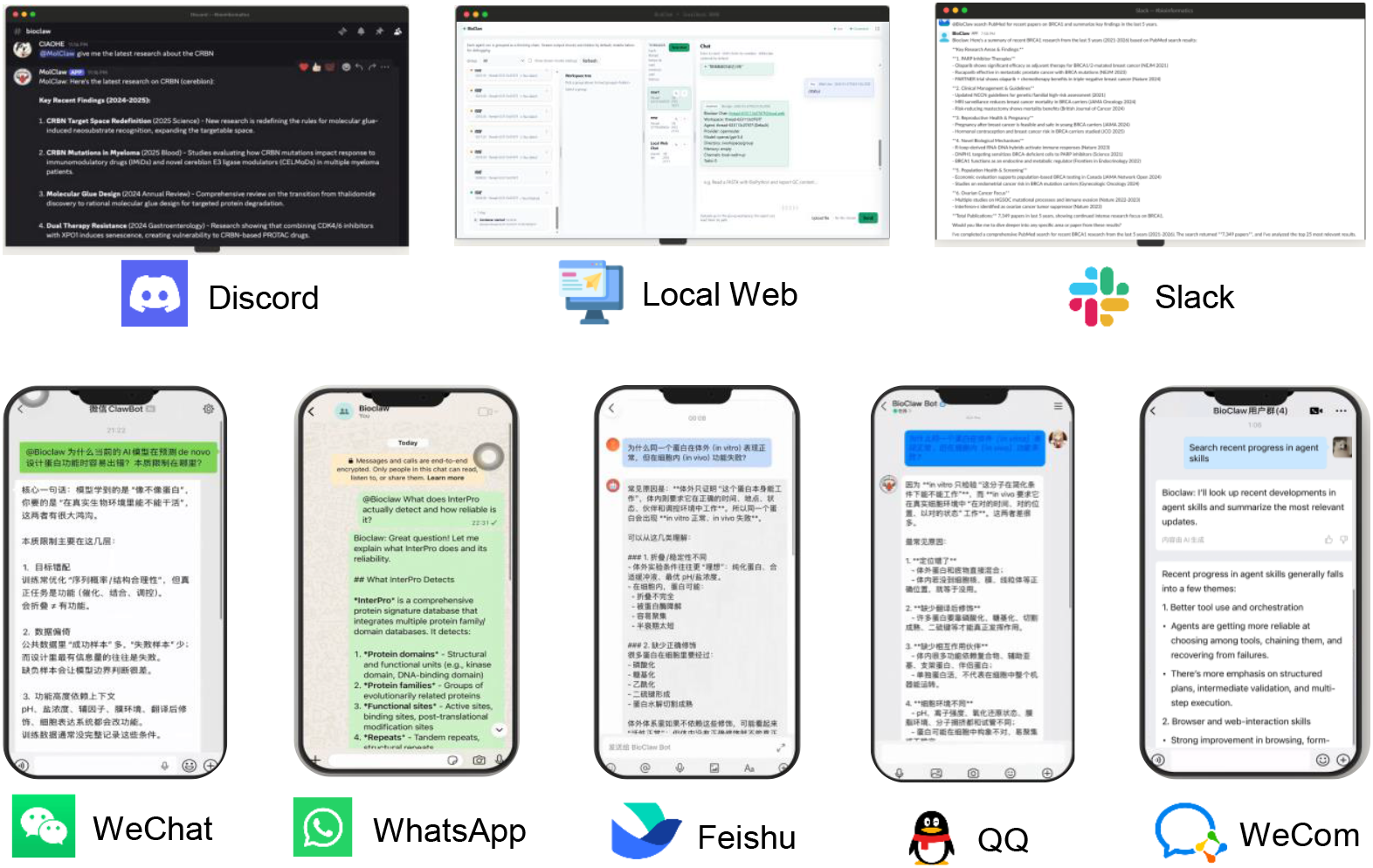
Real-world multi-platform deployment of BioClaw. We implemented and released BioClaw across eight messaging platforms, all backed by a shared orchestration and execution stack, enabling the same bioinformatics capabilities to be accessed from heterogeneous chat environments.

From a systems perspective, this deployment reflects a deliberate architectural contribution: the separation of platform-specific interface layers from a shared backend for orchestration, tool execution, and artifact management. Although user requests originate from different chat platforms, they are routed through the same analytical runtime and therefore inherit the same workspace semantics, tool availability, and artifact-return mechanism. As a result, BioClaw provides consistent bioinformatics functionality across multiple communication ecosystems without requiring platform-specific redesign of the underlying scientific workflow engine.

In addition to the multi-platform deployment, we also developed a public-facing project homepage for BioClaw (Figure 6) to serve as a unified access point for introducing the system, documenting its capabilities, and facilitating public visibility. This homepage complements the messaging-platform deployment by improving discoverability and lowering the barrier to access for new users and collaborators. Together, the cross-platform deployment and the public project page represent practical engineering contributions aimed at making BioClaw usable, reachable, and maintainable beyond a single experimental setting.

**Figure 6.**
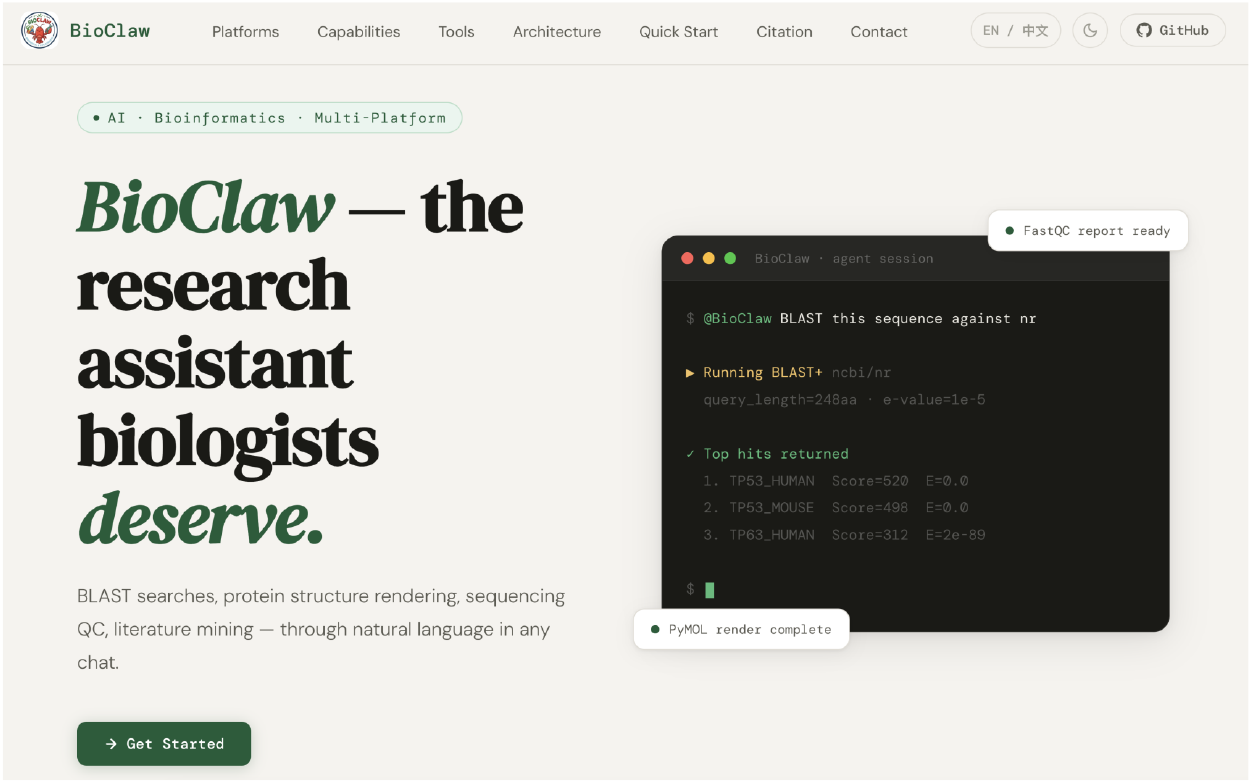
Public home page of BioClaw, developed as part of the project to provide a unified entry point for system introduction, access, and documentation. Available at https://ivegotmagicbean.github.io/BioClaw-Page.

## 5. Related Work

### 5.1. Tool-Using Language Agents

A central line of recent work studies how language models call external tools and environments instead of solving tasks through text-only generation. Early formulations such as Toolformer (Schick et al., 2023) demonstrated self-supervised tool-use learning, while ReAct-style trajectories (Yao et al., 2023) established the value of interleaving reasoning and acting. Subsequent systems focus on stronger execution reliability and trajectory-level tuning (Chen et al., 2023, 2025; Feng et al., 2025). BioClaw belongs to this family but targets a deployment-specific niche: persistent, multi-user chat orchestration with biomedical toolchains and per-group isolation guarantees.

### 5.2. Biomedical AI Agents

STELLA (Jin et al., 2025) highlights a self-evolving, multimodal biomedical agent paradigm where scientific problem solving is grounded in tool creation and iterative refinement. BioClaw is directly inspired by this perspective and translates it into a production-style messaging assistant: users issue requests from chat, the system executes tools in isolated runtimes, and results are returned as conversational artifacts. The two projects are complementary. STELLA primarily advances capability and scientific reasoning methodology, whereas BioClaw emphasizes integration architecture, operations, and reliability in collaborative communication channels.

### 5.3. Containerized Orchestration for Agent Execution

From a systems perspective, BioClaw follows the container-agent orchestration lineage represented by NanoClaw (QwibitAI, 2026) and practical SDK-backed agent runtimes (Anthropic, 2026). Core design patterns include per-tenant runtime isolation, filesystem-mediated interoperability, and persistent metadata for session continuity. Relative to generic orchestrators, BioClaw’s contribution is a domain-adapted execution substrate: pre-bundled bioinformatics tools, group-scoped workspace semantics, and trigger-aware chat routing across multiple channels.

## 6. Discussion

BioClaw demonstrates a practical path from biomedical agent concepts to an operational assistant embedded in team communication channels. By combining channel-agnostic orchestration, per-group container isolation, persistent state management, and a broad bioinformatics runtime, it reduces the friction between scientific discussion and executable analysis. Aligned with STELLA’s tool-centric biomedical agent philosophy, BioClaw further emphasizes deployment properties essential for real-world use, including reliability, recoverability, and secure multi-group operation. Furthermore, the automatic generation of Jupyter notebooks from agent trace events ensures that every conversational analysis session produces a reproducible, re-executable artifact, addressing a key requirement for computational biology workflows. As such, it provides a useful blueprint for conversational and reproducible computational biology assistance.

More broadly, BioClaw suggests that group chat can serve not only as a coordination channel, but also as a medium for scientific discovery. In practice, research ideas often emerge through iterative exchanges: an unexpected result is shared, a hypothesis is proposed, a follow-up analysis is requested, and interpretation is refined collectively. Conventional workflows fragment this process across separate tools for messaging, scripting, visualization, and literature search. By keeping requests, computation, returned artifacts, and interpretation within the same shared thread, BioClaw helps preserve the continuity of collaborative reasoning and shortens the path from discussion to verification.

A promising next step is to extend BioClaw from a single general-purpose assistant to a multi-agent group-chat ecosystem composed of specialized Claws. A single conversation could include, for example, a sequencing-analysis Claw, a structure-biology Claw, a literature-review Claw, a statistics Claw, and a workflow-orchestration Claw, all operating within the same shared context. In such a setting, researchers could discuss a biological question in their existing messaging platform, invoke the most relevant Claw when needed, and combine outputs from multiple specialized agents in one persistent thread. This would bring the division of expertise seen in real scientific teams into the agent layer itself, moving closer to group-chat-native scientific discovery.

## A. Appendix

**Figure S1.**
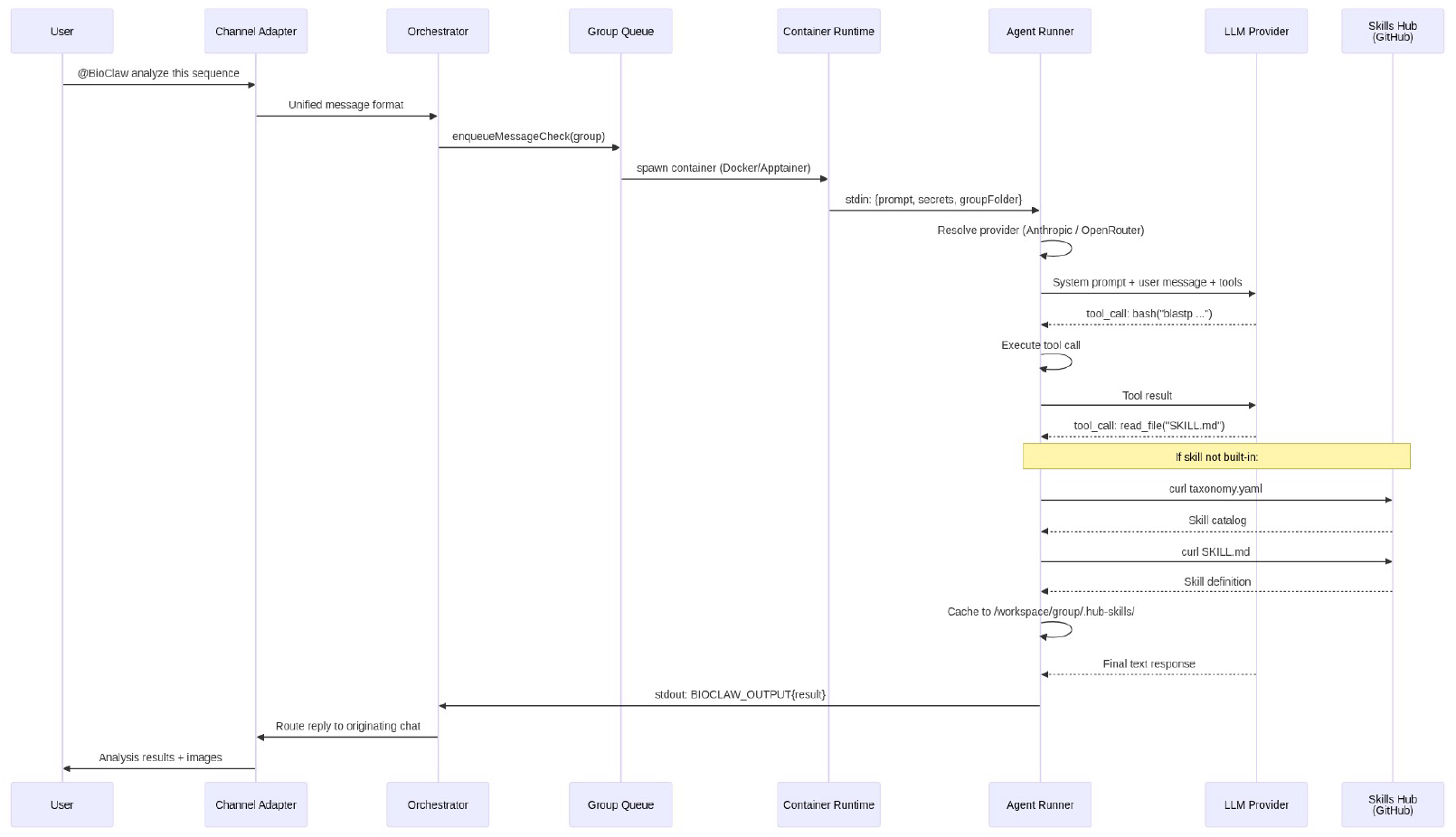
End-to-end execution flow in BioClaw. A user request issued in a chat channel is normalized by the channel adapter and forwarded to the orchestrator, which enqueues the request on a per-group task queue and launches an isolated container through Docker or Apptainer. Inside the container, the agent runner selects the configured model provider, executes tool calls, and accesses built-in skills through local `SKILL.md` files. When a required skill is not built in, the agent dynamically queries the Skills Hub on GitHub, retrieves the corresponding taxonomy entry and skill definition, caches them in the group workspace, and continues execution. Final results are emitted through the stdout sentinel protocol, routed back by the orchestrator, and returned to the originating chat with text and generated images.

**Figure S2.**
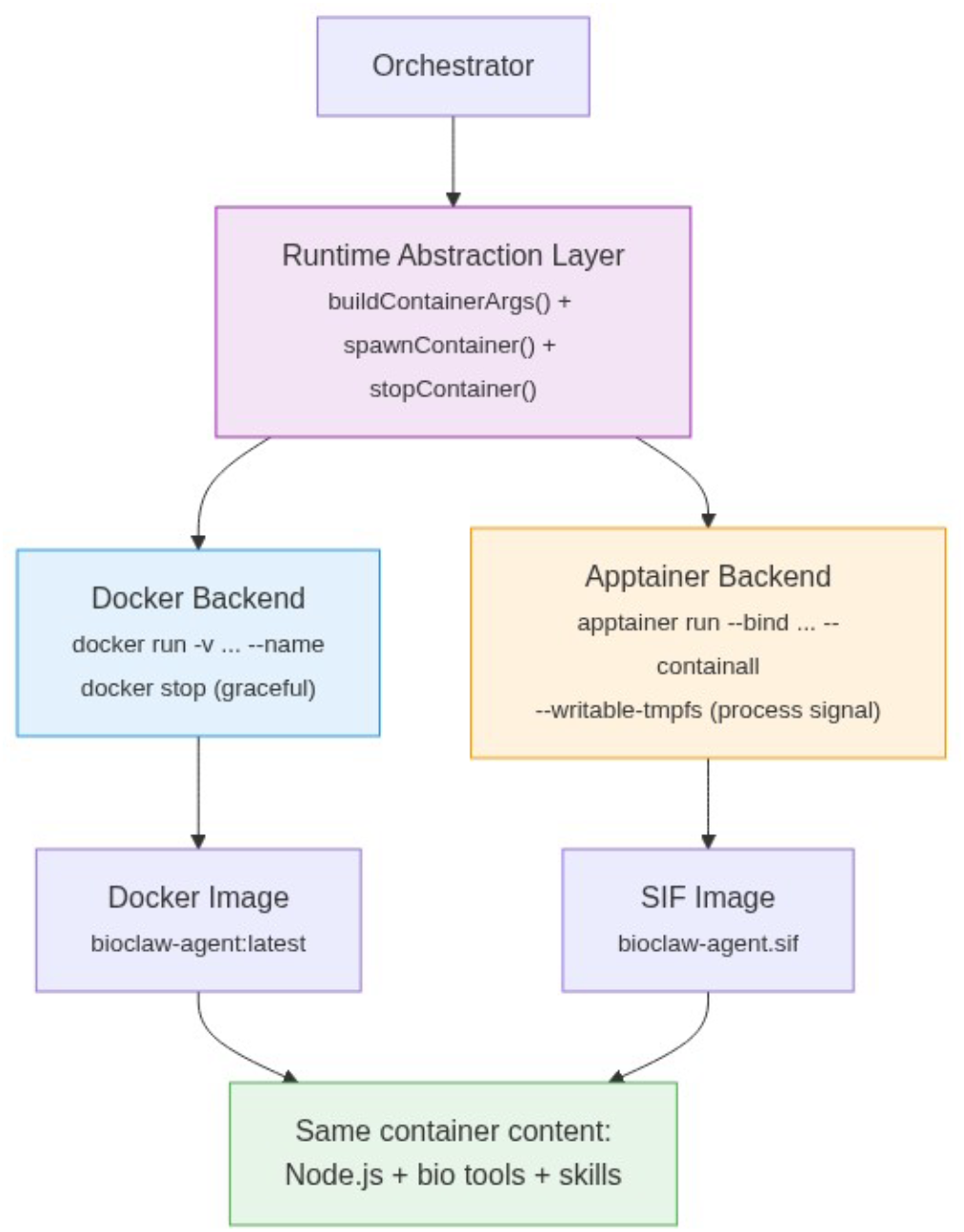
Container runtime abstraction in BioClaw. The orchestrator interacts with a unified runtime layer that supports both Docker and Apptainer backends. While the two runtimes differ in execution mechanics and image format, they provide the same containerized environment for the agent, including Node.js, bioinformatics tools, and skills.

**Figure S3.**
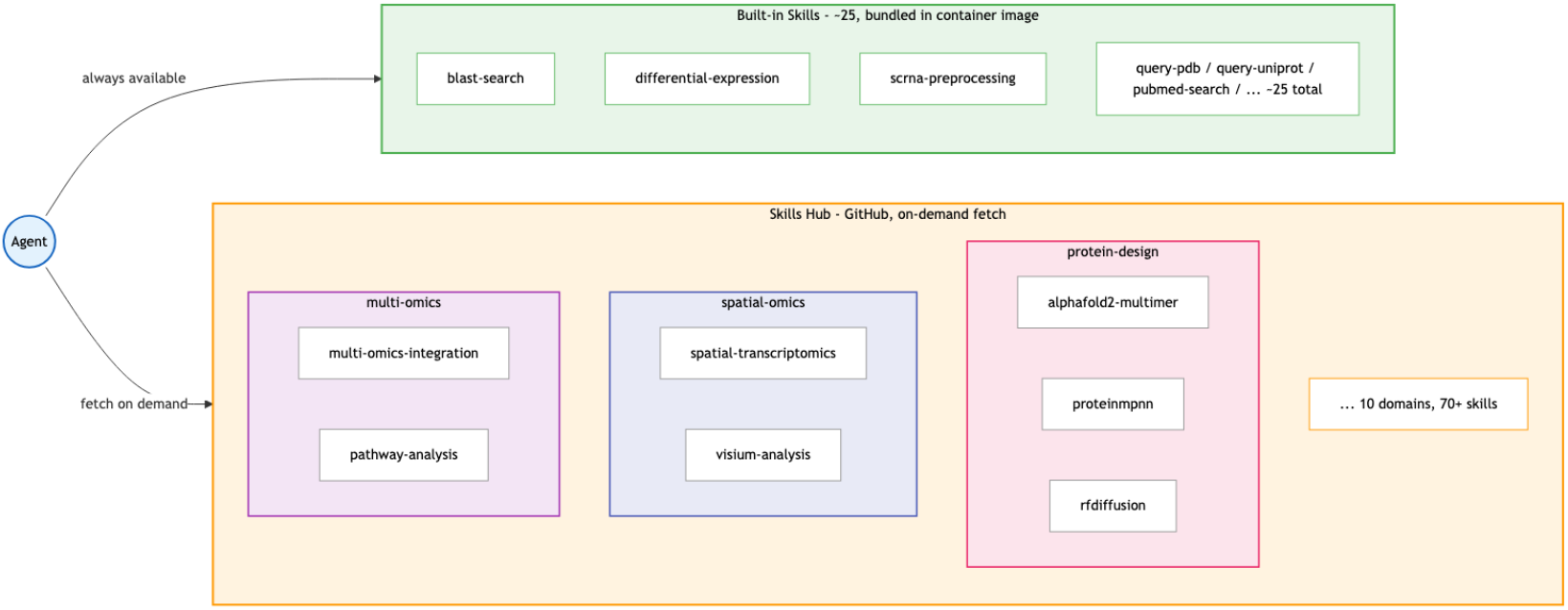
Two-tier skill architecture in BioClaw. The agent has direct access to a set of built-in skills packaged inside the container image, which cover frequently used bioinformatics workflows such as BLAST search, differential expression analysis, scRNA-seq preprocessing, database queries, and literature retrieval. To extend coverage beyond the base image, the agent can also invoke a `skills-hub` meta-skill to discover and fetch additional skills on demand from a community-maintained Skills Hub repository, spanning domains such as multi-omics, spatial omics, and protein design. This design combines low-latency access to core workflows with lightweight, on-demand extensibility to a broader library of specialized analyses.

**Figure S4.**
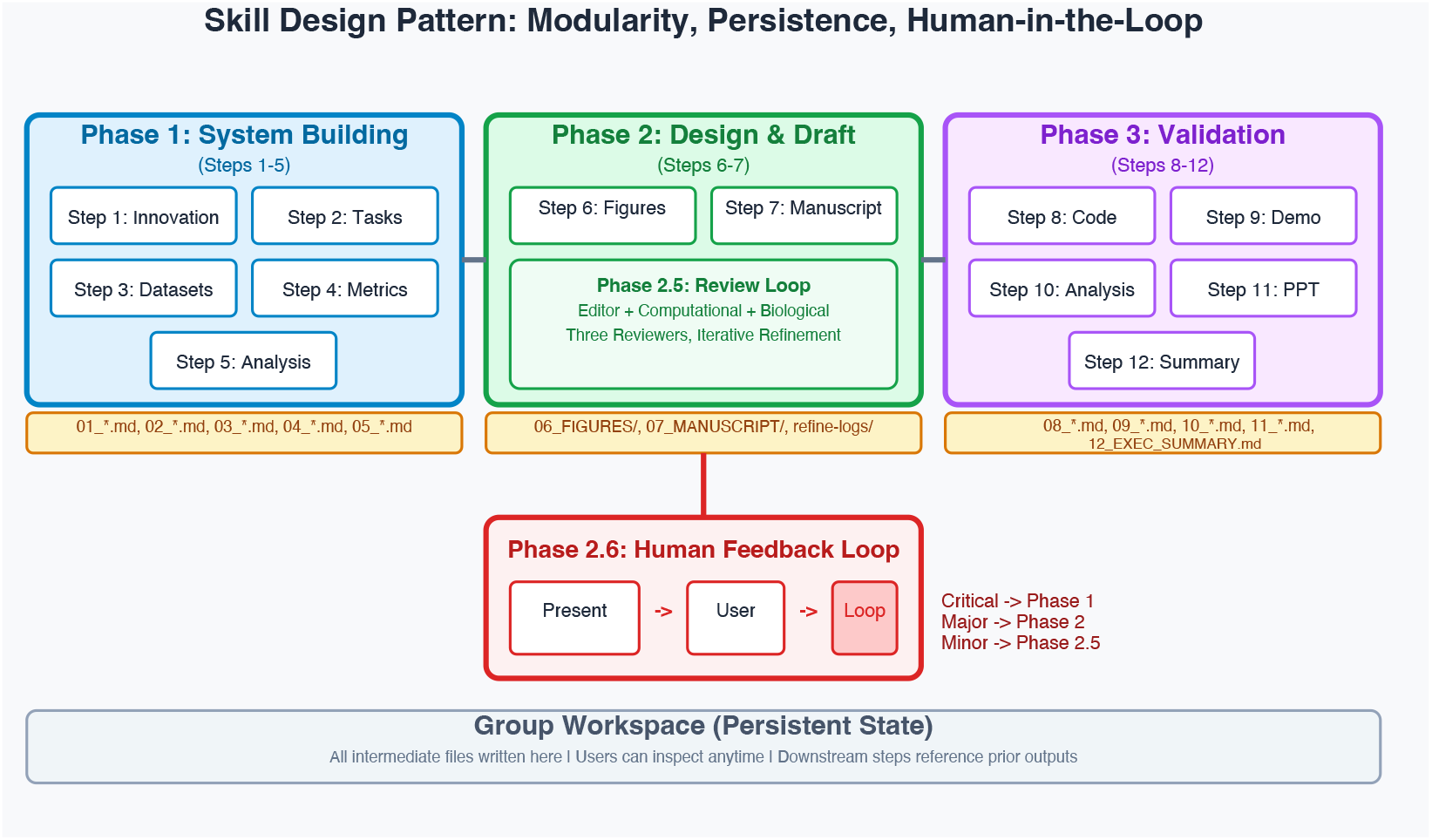
Skill design pattern in BioClaw. A long-horizon Skill is organized into modular phases for system building, design and drafting, and validation, with all intermediate artifacts written to the persistent group workspace. A human feedback loop allows users to inspect results and route revision to the appropriate earlier phase, enabling transparent, resumable, and human-in-the-loop workflow execution.

**Figure S5.**
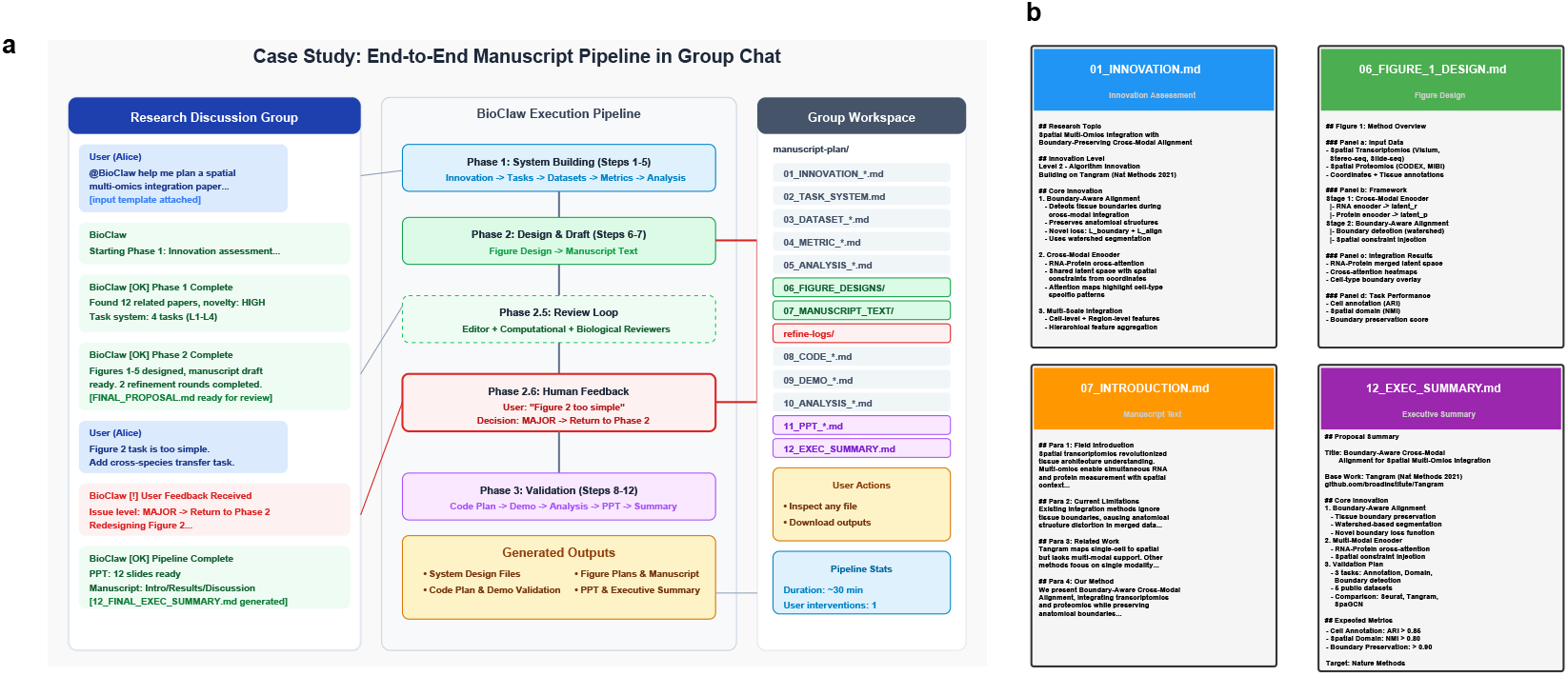
Case study of the end-to-end manuscript-planning Skill in a group chat. (a) A user request is translated into a multi-phase workflow with review and human feedback, while intermediate outputs are continuously materialized in the shared group workspace. (b) Representative generated artifacts from the workflow, including innovation assessment, figure design, manuscript draft, and final execution summary.

**Figure S6.**
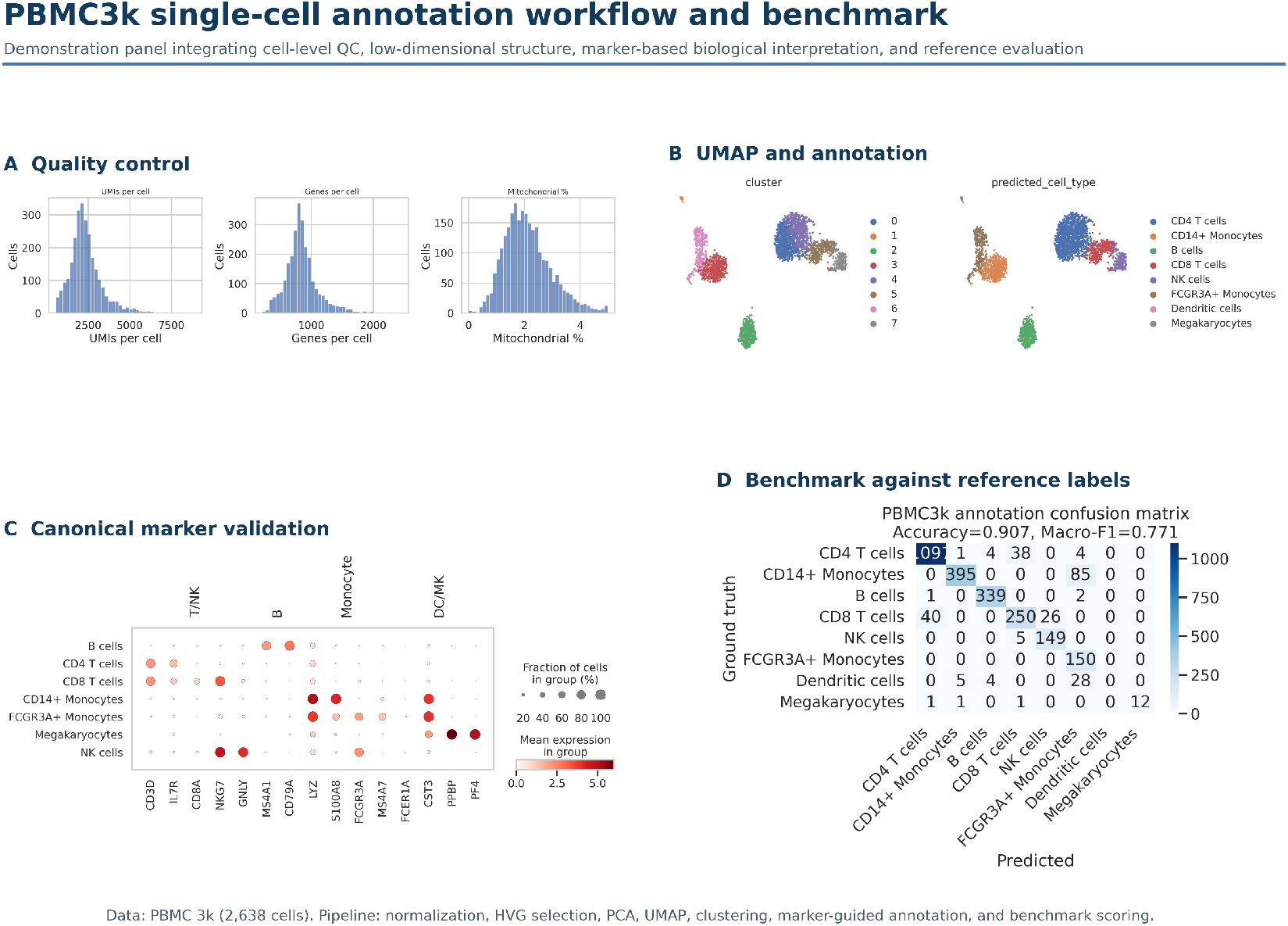
End-to-end single-cell RNA-seq annotation using BioClaw on the PBMC3k dataset. Given an unlabeled single-cell expression matrix (`. h5ad`), BioClaw executes a complete analysis workflow triggered by natural-language instructions, including quality control, normalization, highly variable gene selection, dimensionality reduction (PCA), clustering, and marker-based cell type annotation.(A) Quality-control summaries showing the distributions of UMI counts, detected genes, and mitochondrial transcript percentages (B) UMAP visualization of the processed cells, displaying unsupervised clusters and the corresponding cell types predicted by BioClaw. (C) Dotplot of canonical marker genes across predicted cell populations, supporting biological interpretation of major immune lineages such as T cells, B cells, NK cells, monocytes, dendritic cells, and megakaryocytes. (D) Confusion matrix comparing BioClaw-predicted labels with reference annotations, achieving an accuracy of 0.907 and macro-F1 of 0.771. All steps, from raw data ingestion to visualization and evaluation, are automatically performed by BioClaw within a conversational workflow, demonstrating its ability to translate natural-language requests into executable bioinformatics pipelines with interpretable outputs.

**Figure S7.**
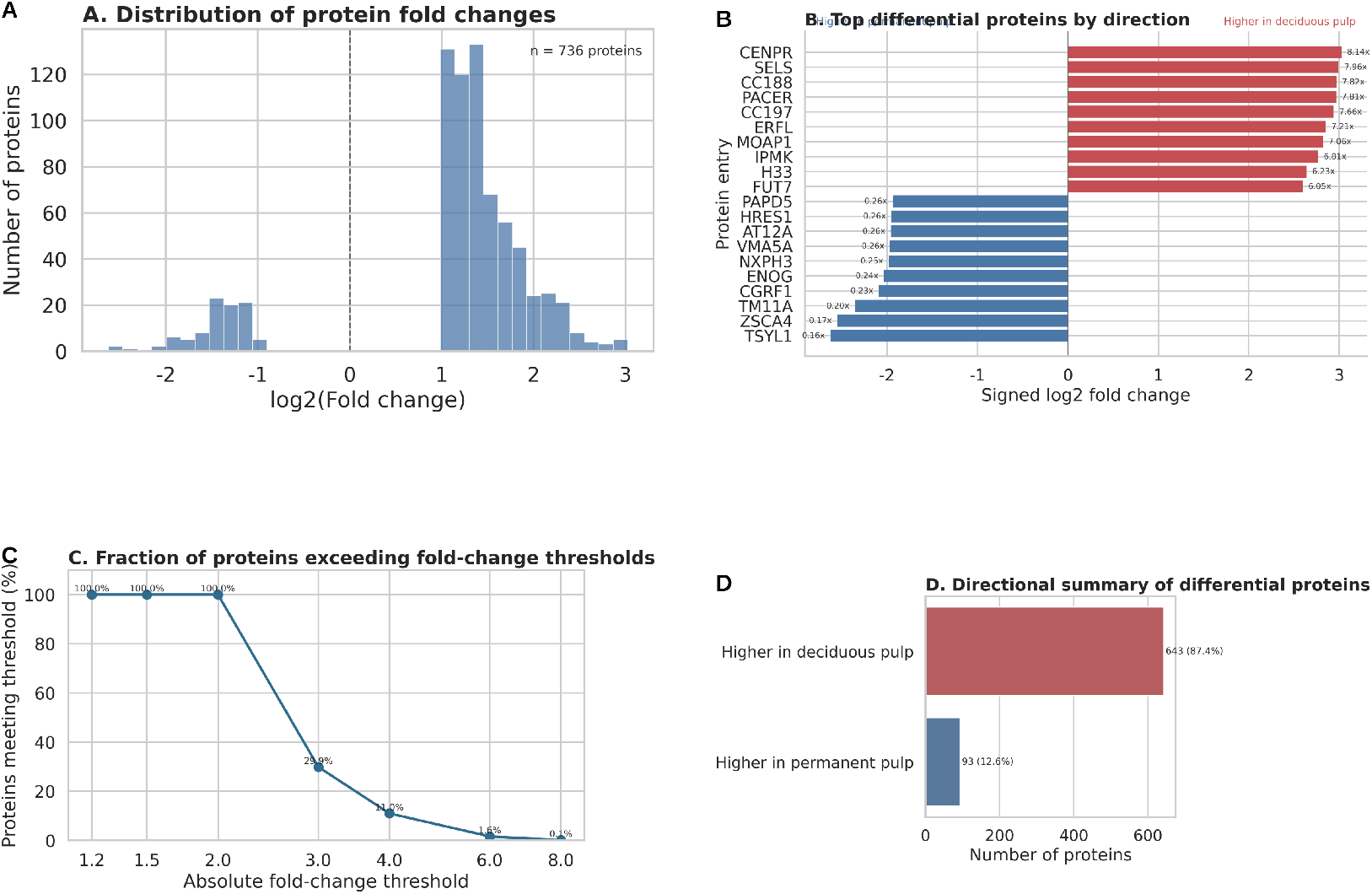
Differential proteomics result summarization using BioClaw. In this task, BioClaw is given an uploaded Excel table containing differential protein results comparing deciduous and permanent dental pulp tissues. The table includes protein identifiers and fold-change values, but does not contain replicate-level quantitative measurements. To analyze this dataset, BioClaw performs a structured interpretation workflow triggered by natural-language instructions. Specifically, it parses the uploaded table, extracts protein-level fold-change values, assigns directionality (fold change > 1: higher in deciduous pulp; fold change < 1: higher in permanent pulp), and organizes the dataset into multiple complementary analytical views instead of re-running statistical testing. (A) Global distribution of log2-transformed fold-change values, providing an overview of effect-size magnitude and directional bias. (B) Representative proteins with the strongest directional abundance shifts, visualized using a mirror bar plot. (C) Fraction of proteins retained under increasingly stringent absolute fold-change thresholds, summarizing the effect-size structure of the dataset. (D) Directional summary showing the number and proportion of proteins in each direction of change. From the uploaded result table, BioClaw extracts 736 differential proteins, of which 643 show higher abundance in deciduous pulp and 93 in permanent pulp, indicating a strongly asymmetric proteomic pattern. All analyses and visualizations are automatically generated by BioClaw from the uploaded result table, demonstrating its ability to interpret heterogeneous biomedical data formats and transform them into structured, presentation-ready analytical summaries within a conversational workflow.

## Footnotes

1 https://github.com/zongtingwei/Bioclaw_Skills_Hub

## Notes

### Competing Interest Statement

The authors have declared no competing interest.

